# Erythropoietin production by the kidney and the liver in response to severe hypoxia evaluated by Western blotting with deglycosylation

**DOI:** 10.1101/2020.03.11.987586

**Authors:** Yukiko Yasuoka, Takashi Fukuyama, Yuichiro Izumi, Yushi Nakayama, Hideki Inoue, Kengo Yanagita, Tomomi Oshima, Taiga Yamazaki, Takayuki Uematsu, Noritada Kobayashi, Yoshitaka Shimada, Yasushi Nagaba, Masashi Mukoyama, Tetsuro Yamashita, Yuichi Sato, Katsumasa Kawahara, Hiroshi Nonoguchi

**Author notes:** **Corresponding author** Hiroshi Nonoguchi, M.D., Ph.D., Division of Internal Medicine, Kitasato University Medical Center, 6-100 Arai, Kitamoto, Saitama 364-8501, Japan.

## Abstract

The detection of erythropoietin (Epo) protein by Western blotting has required pre-purification of the sample. We developed a new Western blot method to detect plasma and urinary Epo using deglycosylation. Epo in urine and tissue and erythropoiesis-stimulating agents (ESAs) in urine were directly detected by our Western blotting. Plasma Epo and ESAs were detected by our Western blotting after deglycosylation. The broad bands of Epo and ESAs were shifted to 22 kDa by deglycosylation except PEG-bound epoetin β pegol. The 22 kDa band from anemic patient urine was confirmed by Liquid Chromatography/Mass Spectrometry (LC/MS) to contain human Epo.

Sever hypoxia (7% O_2,_ 4 hr) caused a 400-fold increase in deglycosylated Epo expression in rat kidneys, which is consistent with the increases in both Epo gene expression and plasma Epo concentration. Immunohistochemistry showed Epo expression in nephrons but not in interstitial cells under control conditions, and hypoxia increased Epo expression in interstitial cells but not in tubules.

These data show that intrinsic Epo and all ESAs can be detected by Western blot either directly in urine or after deglycosylation in blood, and that the kidney is the main and sole site of Epo production in control and severe hypoxia. Our method will completely change Epo doping and detection.

## Introduction

Anemia is one of the most common diseases in humans [1]. Severe anemia and hypoxia stimulate the production of erythropoietin (Epo) by the kidney [2-8]. The increase in Epo production is measured by the increases in serum and urine Epo concentrations and in Epo mRNA expression in the kidney [4-11]. Serum or urine Epo concentrations have been measured by radio immunoassay (RIA) or enzyme-linked immunosorbent assay (ELISA) using antibodies against Epo [4, 9-12]. However, Epo protein expression in the kidney or liver has not been measured accurately, since Western blotting of Epo has not been possible. Serum/urine Epo concentrations, and Epo mRNA and HIF1α/2α expressions in the kidney and liver have been used as a substitute for Epo protein expression in the kidney and liver [4-13]. However, the increase of kidney-produced Epo has not been shown to increase to the same degree. This suggest the possibility that Epo production by the liver may have some role for the increase of Epo production in response to severe hypoxia [2, 3, 14].

The discovery of Epo led to the invention of erythropoiesis stimulating agents (ESAs) to treat anemic patients with chronic kidney disease (CKD) [15-17]. ESAs have also been illegally used by athletes to improve physical activity, leading to tests for doping [18]. The World Anti-Doping Agency (WADA) Technical Documents for Epo (TD2014EPO in TD2019INDEX) recommended the use of isoelectrical focusing (IEF) and/or SAR-PAGE after enrichment for Epo through ultrafiltration, selective protein precipitation or immunopurification to detect Epo in the urine or serum/plasma [19]. ELISA or Liquid Chromatography/Mass Spectrometry (LC/MS) after the pre-purification of urine are also useful. These recommendations clearly show that the detection of Epo by Western blotting is difficult.

We have reported a new method of Western blot analysis succeeding in the detection of kidney-produced Epo [20]. We have reported that Epo is produced by the cortical nephrons in control condition using in situ hybridization, immunohistochemistry and real time PCR with microdissected nephron segments. We also showed that Epo production by the intercalated cells of the collecting ducts is regulated by renin-angiotensin-aldosterone system [20]. We modified our method to detect plasma and urinary Epo. We report the new Western blot method for the detection of Epo protein in the plasma or urine. Using our new method, we investigated the role of kidney and liver for Epo production in response to severe hypoxia.

## Methods

### Materials and animals

Male Sprague Dawley rats (Japan SLC, Hamamatsu, Japan) were used in our study. In the severe hypoxia experiments, rats were exposed to 7% O_2_ and 93% N_2_ for 1-4 hr, which is known to stimulate rapid Epo production and is closer to the conditions at the summit of Mount Everest [9, 21]. For the detection of ESAs in plasma and urine, large doses of ESAs were administered to some rats through the vena cava, and plasma and urine were collected after 30 min from the aorta and bladder, respectively. Animal experiments were conducted in accordance with the Kitasato University Guide for the Care and Use of Laboratory Animals and were approved by the Institutional Animal Care and Use Committee (Approval No. 2018-030, 25-2). Blood and urine were collected from patients with CKD who received ESAs and from patients with severe anaemia. Urine was concentrated using a Vivaspin (GE Healthcare Bio-Science AB, Sweden). Our protocols were checked and approved by the above committee and the Ethics Committee at Kitasato University Medical Center (25-2, 2018032, 2019029). Informed consent was obtained from all patients.

### Real-time PCR in control and hypoxic rats

The renal cortex and liver were collected from control and hypoxic rats. RNA was extracted using the RNeasy Mini Kit (Qiagen, 74106) and Qiacube. cDNA was synthesized using a Takara PrimeScript™ II 1st strand cDNA Synthesis Kit (Takara, 6210). Real-time PCR was performed using probes from Applied Biosystems and Premix Ex Taq (Takara, RR39LR). Probes were obtained from Applied Biosystems (Epo, Rn01481376_m1; HIF2α, Rn00576515_m1; HIF1α, Rn01472831_m1; PHD2, Rn00710295_m1, Thermo Fisher Scientific, USA). β-actin (Rn00667869_m1) was used as an internal standard.

### Western blot analysis

Western blot analysis was performed as described previously [20, 22]. Protein was collected from the renal cortex and liver using CelLytic MT (Sigma-Aldrich, C-3228) plus protease inhibitor (Roche, 05892970001). Urine samples were obtained from rats injected large doses of ESAs 30 min before the collection and from anemic patients. Plasma was obtained from rats injected large amount of ESAs and from patients with iron deficiency anaemia or CKD. An anemic patient was treated by iron supplementation and blood was collected at severe and mild anemia and after complete recovery. Blood was also collected form CKD patients who were treated by the injection of epoetin β pegol and control subject. Informed consent was obtained from all patients. Urine samples were concentrated by Vivaspin (GE Healthcare Bio-Science AB) and used for western blot. Plasma samples were used directly or after deglycosylation as described below. After SDS-PAGE, proteins were transferred to a PVDF membrane (Immobilon-P, Merck Millipore, IPVH00010) with 160 mA for 90 min. The membrane was blocked with 5% skim milk (Morinaga, Japan) for 60 min and incubated with the antibody against Epo (Santa Cruz, sc-5290, 1:500-2,000) for 60 min at room temperature. After washing, the membrane was incubated with a secondary antibody (the goat anti-mouse IgG (H+L) (Jackson ImmunoResearch Laboratories, 115-035-166, 1:5,000) for 60 min. Bands were visualized by the ECL Select Western Blotting Detection System (GE Healthcare Bio-Science AB, RPN2235) and LAS 4000 (Fujifilm). The band intensity was normalized against that of β-actin (MBL, M177-3), which was measured after stripping and reprobing the membrane (stripping solution, Wako, RR39LR). In some experiments, another antibody against Epo (clone AE7A5, MAB2871, R & D Systems) was used to compare the specificity of the antibody.

### Deglycosylation study

Since the Epo protein is a glycosylated protein, deglycosylation was performed. N-glycosidase F (PNGase, Takara, 4450) was used as previously reported [22]. In brief, a mixture of 7.5 μl of plasma, 2.5 μl of water, and 1μl of 10% SDS was boiled for 3 min. Then, 11 μl of 2x stabilizing buffer was added, and 2 μl of PBS or PNGase was added. The samples were incubated in a water bath at 37°C for 15-20 hr. After incubation, the samples were spun down, and the supernatant was collected. For urine analysis, 7.5 μl – 30 ml of urine was used either directly or after concentration by Vivaspin. To 10 μl of concentrated urine, 1 μl of 10% SDS was added and boiled for 3min. The subsequent steps were the same as those performed for plasma. In the kidney and liver samples, 10 μl samples were treated in the same manner as urine. The 2x stabilizing buffer contained 62.5 mM Tris-HCl (pH 8.6), 24 mM EDTA, 2% NP-40 and 4% 2-mercaptoethanol.

### Plasma Epo concentration measurements

Plasma and urine were collected from control and hypoxic rats. Plasma, serum and urine were also collected from patients with renal anaemia treated with ESAs or from patients with iron-deficient anaemia. Plasma, serum and urine Epo concentrations were measured by CLEIA (SRL, Tokyo, Japan, using Access Epo by Beckman Coulter, Brea, USA).

### Immunohistochemistry of Epo production sites

Immunohistochemistry (IHC) of Epo expression was performed in control and severe hypoxic rats as previously reported [20, 23, 24]. A polyclonal antibody against the same sequences as sc-5290 was used, namely, sc-7956. Images were obtained using an optical microscope (Axio Imager. M2, Carl Zeiss, Oberkochen, Germany) with a digital camera (AxioCam 506, Carl Zeiss). Captured images were analysed using an image analysis system (ZEN 2, Carl Zeiss).

### LC/MS analysis of band from western blot

The 22 kDa band of the western blot was excised and subjected to LC/MS as previously reported [25]. Negative staining was used to detect deglycosylated recombinant Epo. The negatively stained protein bands were excised from the SDS-PAGE gel, and in-gel tryptic digestion was carried out using ProteaseMAX reagent (Promega, WI, USA) according to the manufacturer’s protocol. The peptides were separated by L-column2 ODS (3 μm, 0.1 x 150 mm, CERI, Tokyo, Japan) at a flow rate of 500 nl/min using a linear gradient of acetonitrile (5% to 45%). Nano-LC-MS/MS analyses were performed with an LTQ-Orbitrap XL mass spectrometer (Thermo Fisher Scientific, MA, USA) as previously described [25].

### Statistical analyses

Statistical analyses were performed using Excel Statics (BellCurve, Tokyo, Japan). Statistical significance was analysed using ANOVA and multiple comparison with Dunnett test, or non-parametric analysis by the Kruskal-Wallis test and multiple comparisons by the Shirley-Williams test. P<0.05 was considered statistically significant.

## Results

### Detection of Epo protein

We have reported that our western blot recognized hypoxic rat kidney Epo protein and the deglycosylated protein at 34-43 and 22 kDa, respectively. The specificity of sc-5290 was better than that of AE7A5 (Fig. 1A, B). ESAs were also detected by Western blot, and deglycosylation caused a shift of the bands to 22 kDa, except for that of epoetin β pegol (Fig. 2A1, A2). The deglycosylated band at 22 kDa showed a 10-100 times lower limit of detection than the non-deglycosylated band at 34-43 kDa (Fig. 2B1, 2B2).

**Fig. 1.**
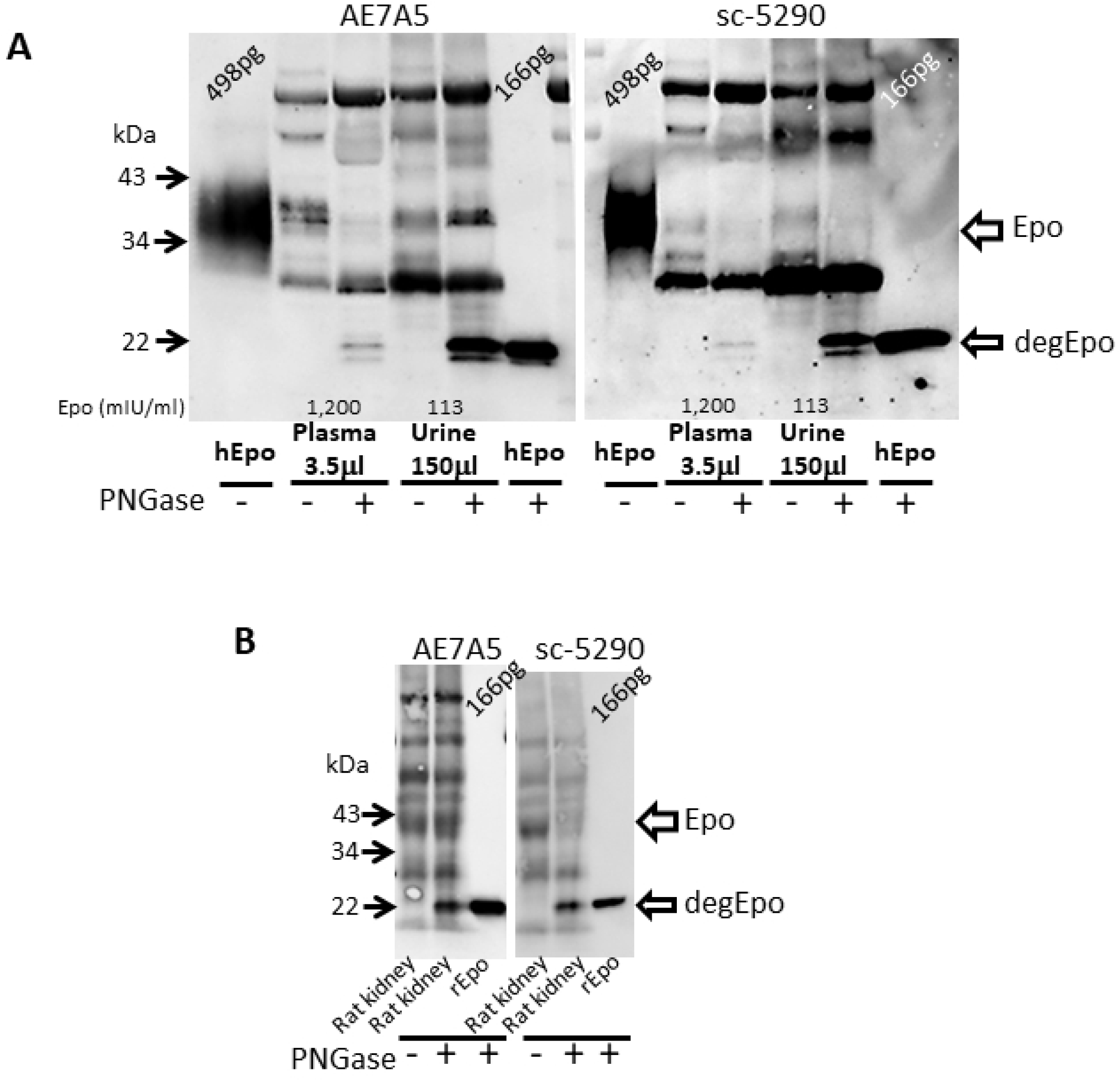
Comparison of AE7A5 and sc-5290. **A.** Plasma and concentrated urine from anemic patients were used for western blotting with or without deglycosylation. Although both AE7A5 and sc-5290 recognize Epo at 34-43 and 22 kDa, the specificity of sc-5290 was better than that of AE7A5 especially after deglycosylation. **B.** The kidney cortex from hypoxic rats were used for western blotting. Although a 34-43 kDa band by sc-5290 became pale after deglycosylation, same band by AE7A5 remains strong after deglycosylation. hEpo; recombinant human Epo, rEpo; recombinant rat Epo.

**Fig.2.**
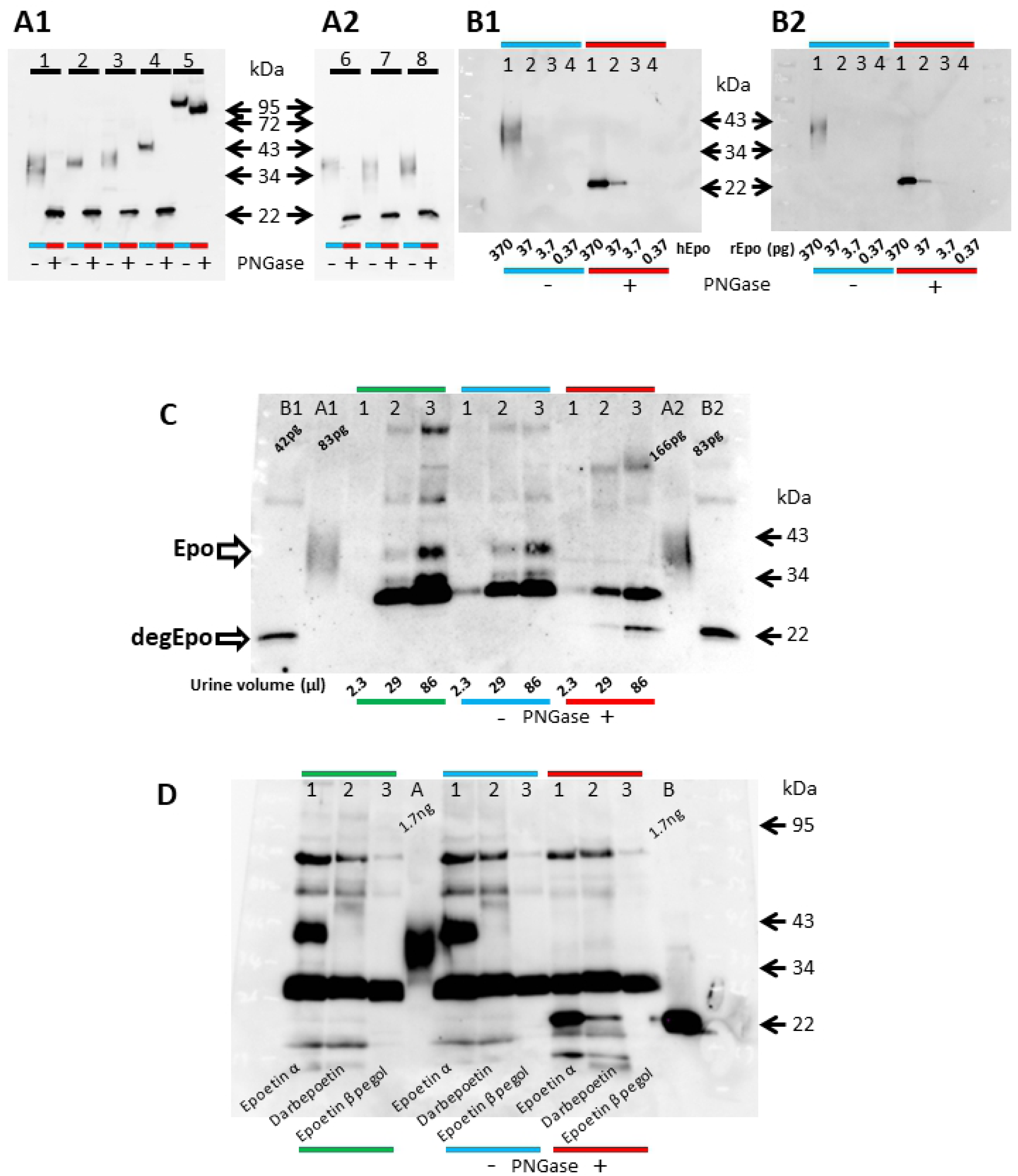
Detection of Epo and ESAs in urine by western blotting. **A1.** Expression of recombinant human Epo and ESAs detected by Western blotting. Recombinant human Epo shows a broad band at 34-43 kDa. Epoetin α and β, darbepoetin and epoetin β pegol gradually increased in size. Deglycosylation shifted all human Epo and ESAs to 22 kDa except PEG-bound epoetin β pegol. Lane 1: recombinant human Epo; Lane 2, epoetin α; lane 3, epoetin β; lane 4, darbepoetin; and lane 5, epoetin β pegol. The left and right lanes of each peptide are without and with deglycosylation, respectively. **A2.** Expression of rat (lane 6), mouse (lane 7) and human Epo (lane 8). Rat, mouse and human Epo showed the same expression at 34-43 kDa, and deglycosylation shifted all bands to 22 kDa. **B1.** Expression of recombinant human Epo in glycosylated (blue line) and deglycosylated forms (red line). The detection limits of glycosylated and deglycosylated human Epo were 370 and 37 pg, respectively. **B2.** The detection limit of glycosylated and deglycosylaed recombinant rat Epo was 370 and 3.7 pg, respectively. **C.** Detection of intrinsic Epo in human urine. Urine from anemic patient was applied to the western blot: 2.3, 29 and 86 μl samples of urine (Epo concentration 152 mIU/ml) were concentrated by Vivaspin and used in lanes 1, 2 and 3, respectively. Deglycosylated Epo was observed in more than 29 μl of urine. A, B: glycosylated and deglycosylated recombinant human Epo, respectively. Green lien, direct application; blue line, incubation with deglycosylation buffer; and red line, after deglycosylation. **D.** Detection of ESAs in rat urine. Male SD rats (200 g) were injected with epoetin α (600 μg), darbepoetin (4.5 μg) and epoetin β pegol (3.8 μg), and urine was obtained after 30 min. The plasma Epo concentrations of each rat were 37,800, 29,400 and 527 mIU/ml for epoetin α, darbepoetin and epoetin β pegol, respectively. The direct analysis of urine (5 μl) showed a clear and broad band of epoetin α (sample 1) at 34-43 kDa. The band of darbepoetin (sample 2) was pale and that of epoetin β pegol (sample 3) was not observed. The band of darbepoetin became slightly clearer after the incubation of urine with deglycosylation buffer (blue line). The bands of epoetin α and darbepoetin were shifted to 22 kDa. The deglycosylated band of darbepoetin (sample 2 in red line) was clearer than the glycosylated band. Since the rat urine samples were very small, the urine Epo concentration was not measured. A and B: glycosylated and deglycosylated rat Epo, respectively.

### Detection of Epo protein and ESAs in urine

The direct analysis (green line) and incubation with deglycosylation buffer (blue line) of anemic patient’s urine both volume-dependently showed an Epo protein band at 36-40 kDa. Deglycosylation (red line) shifted the bands to 22 kDa (Fig. 2C). Epoetin α (lane 1) and darbepoetin (lane 2) were detected by the direct application of rat urine after bolus injection. Epoetin β pegol (lane 3) was not detected, probably due to its limited excretion into the urine (Fig. 2D).

### Detection of Epo protein and ESAs in plasma

The direct analysis of plasma from control and hypoxic rats by Western blotting showed no band (Fig. 3A, green line). Incubation of the plasma with deglycosylation buffer showed bands at 34-43 kDa in 4-hr hypoxic rats but not in control rats (Fig. 3A, blue line). Deglycosylation shifted the broad band at 34-43 kDa to 22 kDa (Fig. 3A, red line). Next, direct analysis of plasma from anemic patient also showed no band (Fig. 3B, green line). Incubation of the plasma with deglycosylation buffer showed a broad band at 36-40 kDa only in the case of severe anemia (Fig. 3B, lane 1, blue line). The partial recovery of anemia caused a faint band at 36-40 kDa, and complete recovery revealed no broad band at approximately 36-40 kDa. Deglycosylation caused an intense band at 22 kDa in anemia, and partial recovery of anemia caused a very faint band at 22 kDa (Fig. 3B, red line). No band was observed at 22 kDa after complete recovery.

**Fig.3.**
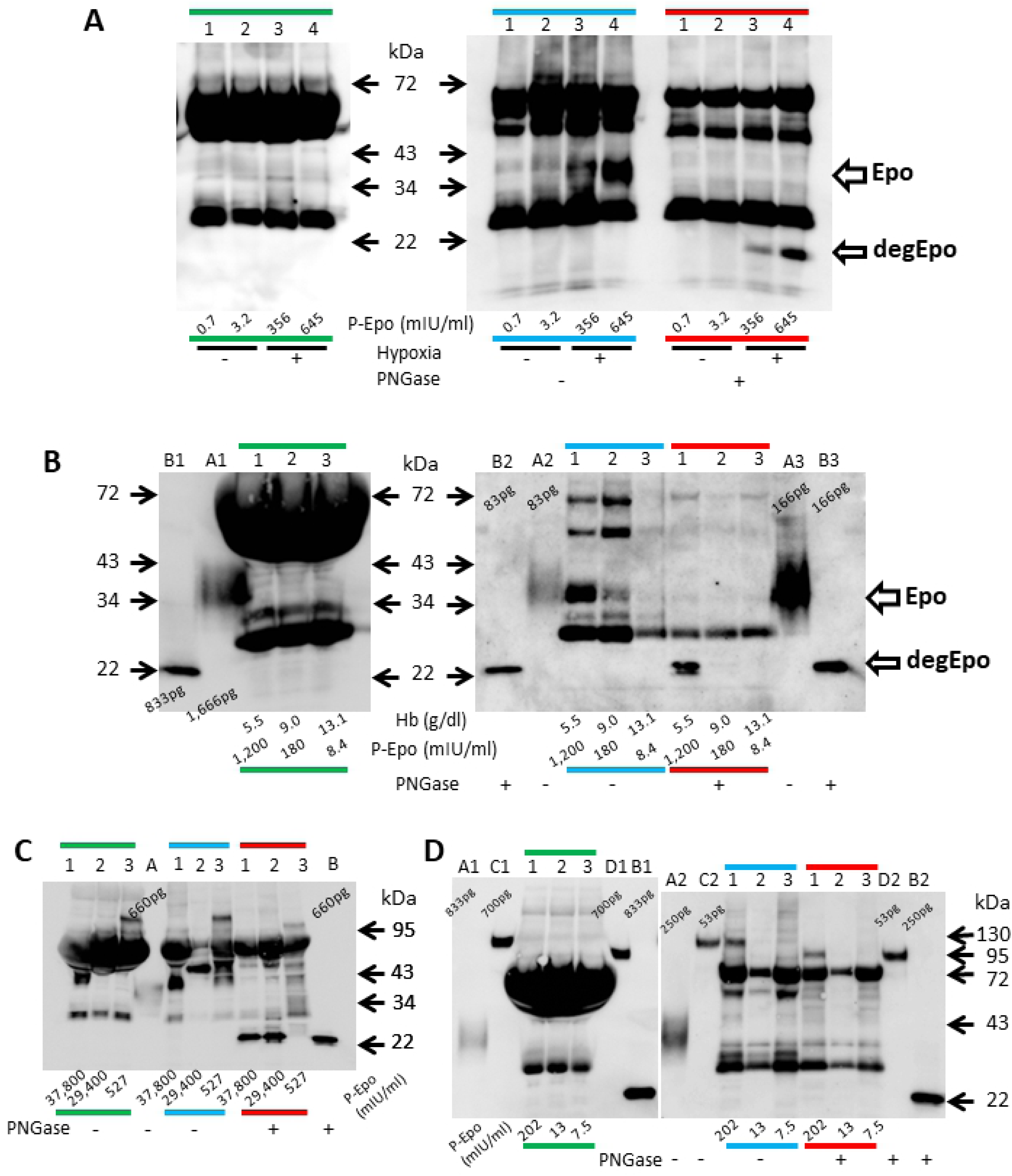
Detection of Epo and ESAs in plasma. **A.** Detection of intrinsic rat Epo in control and hypoxic rats. Although no bands were observed by in the direct analysis of plasma (2 μl) (green line), the incubation of plasma from hypoxic rats (7 μl) with deglycosylation buffer (blue line) resulted in the appearance of Epo bands at 34-43 kDa, which were shifted to 22 kDa by deglycosylation (red line). Lanes 1-2, control rats. Lanes 3-4, hypoxic rats. The plasma Epo concentrations in each rat were 0.7, 3.2, 356 and 645 mIU/ml, respectively. The green, blue and red lines show direct application and incubation with deglycosylation buffer without and with PNGase, respectively. **B.** Detection of intrinsic human Epo in the plasma of a patient with severe anemia. Plasma was obtained under severe and mild anaemia and after recovery (plasma haemoglobin levels were 5.5, 9.0, and 13.1 g/dl, respectively). No bands were observed with the direct analysis of plasma (2 μl). Incubation of plasma (5 μl) with deglycosylation buffer revealed the band at 36-40 kDa only under anemic conditions, and the bands were shifted to 22 kDa. Plasma Epo concentrations were 1,200, 180, and 8.4 mIU/ml, respectively. A, B; glycosylated and deglycosylated recombinant human Epo, respectively. **C.** Detection of ESAs in rats injected with a large doses of ESAs. Male SD rats were injected with epoetin α, darbepoetin and epoetin β pegol as described in Fig. 2D, and blood was obtained after 30 min. The bands of epoetin α and epoetin β pegol were observed by the direct analysis of plasma (2 μl), while the band of darbepoetin was obscured by a non-specific band (green line). Incubation of plasma (5 μl) with deglycosylation buffer reduced the non-specific band, and the band of darbepoetin became clear (blue line). The bands of epoetin α and darbepoetin shifted to 22 kDa, while the band of epoetin β pegol was slightly reduced in size (red line). A, B; glycosylated and deglycosylated recombinant rat Epo, respectively. **D.** Detection of plasma ESA in patients. Plasma samples from patients treated with epoetin β pegol and control subjects were subjected to western blotting. No bands were observed by the direct analysis of plasma (2 μl) (green line). The incubation of plasma (3.5 μl) with deglycosylated buffer revealed the band corresponding to epoetin β pegol in patient 1 at 95-130 kDa (blue line). The band was shifted to 80-95 kDa by deglycosylation (red line). The plasma Epo concentrations of each subject were 202, 13, and 7.5 mIU/ml, respectively. Patient 1: serum creatinine 11.93 mg/dl, Hb 8.2, epoetin β pegol injection 3 days before. Patient 2: serum creatinine 3.15 mg/dl, Hb 10.8 g/dl, epoetin β pegol injection 28 days before. Patient 3: serum creatinine 0.73 mg/dl, Hb 15.1 g/dl, no injection. A, B: glycosylated and deglycosylated recombinant human Epo, respectively. C, D: glycosylated and deglycosylated epoetin β pegol, respectively.

The detection of ESAs in plasma was tested in rats after the intravenous injection of large doses of ESAs. The plasma Epo concentration was more than 100 times higher than under severe hypoxia. In this condition, epoetin α and epoetin β pegol were detected by the direct analysis of plasma (Fig. 3C, green line). The band of darbepoetin overlapped with the non-specific band, which was removed by the incubation of plasma with deglycosylation buffer (Fig. 3C, blue line). The bands of epoetin α and darbepoetin were shifted to 22 kDa by deglycosylation (Fig. 3C, red line). The band of epoetin β pegol shifted from 95-120 to 80-95 kDa. In contrast, no band representing epoetin β pegol was detected by the direct analysis of plasma from anemic CKD patients (Fig. 3D, green line). The incubation of plasma with deglycosylation buffer induced the appearance of a band at 95-120 kDa (Fig. 3D, lane 1 in blue line), which was shifted to 80-95 kDa by deglycosylation (Fig. 3D, lane 1 in red line).

### Detection of Epo protein by LC/MS

To confirm that the band at 22 kDa is Epo protein, the 22 kDa band of recombinant human Epo and anemic patient’s urine were excised and analysed by LC/MS (Fig. 4A, B). Seven, and three peptide sequences of human Epo protein (sequence coverage 20% and 12%) were identified in the sample of recombinant human Epo and anemic patient, respectively (Table. 1). Recombinant rat Epo was also identified by LC/MS (Table. 1).

**Table. 1.**
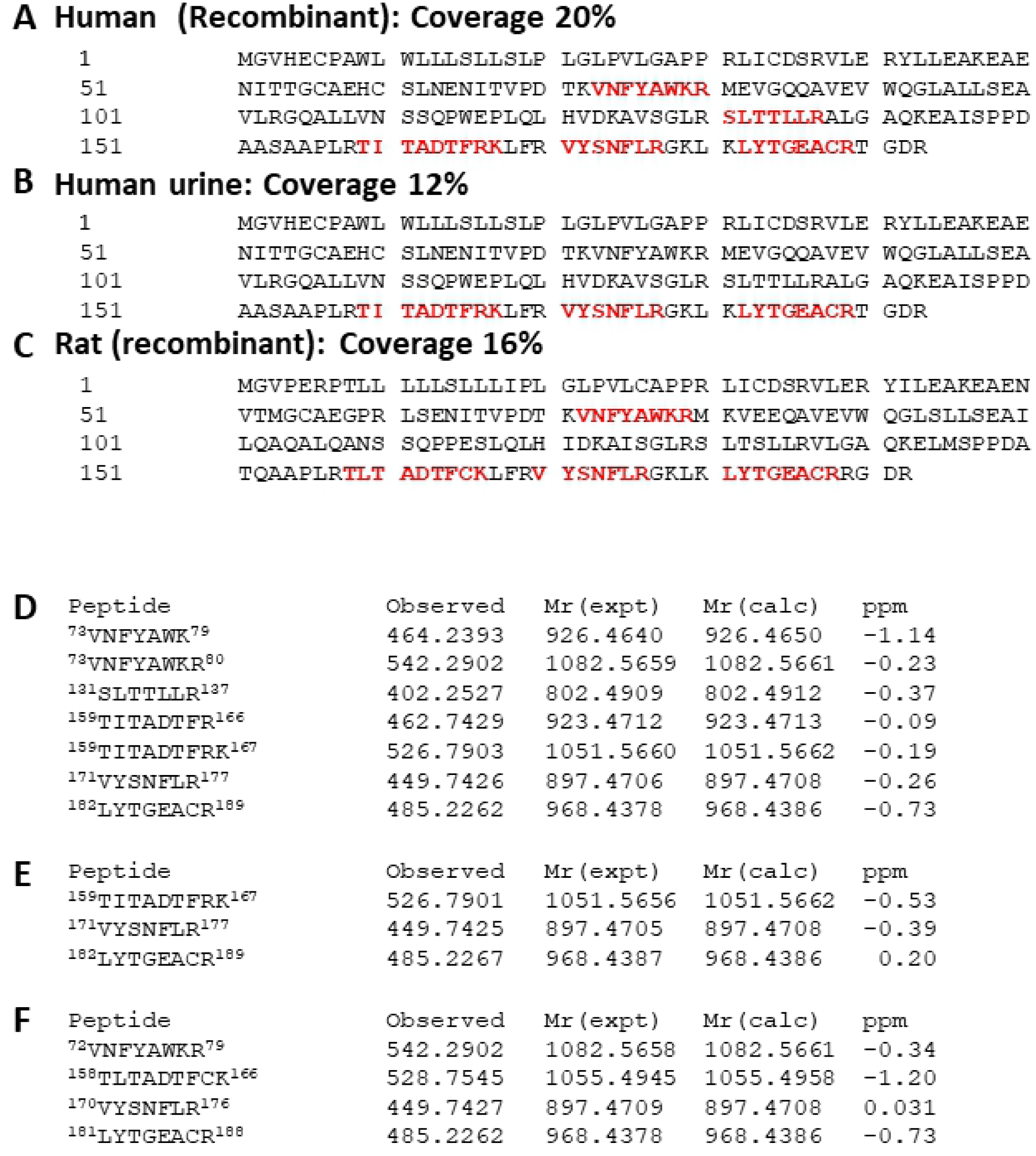
A-C. LC/MS analysis of the 22 kDa band of recombinant human Epo (8.3 ng), concentrated human urine from anaemic patients and recombinant rat Epo (4.4 pg), respectively. Matched peptides are shown in bold red. **D-F.** Detailed LC/MS data on matched peptides of recombinant human Epo (D), human urine sample (E) and recombinant rat Epo (F).

**Fig. 4.**
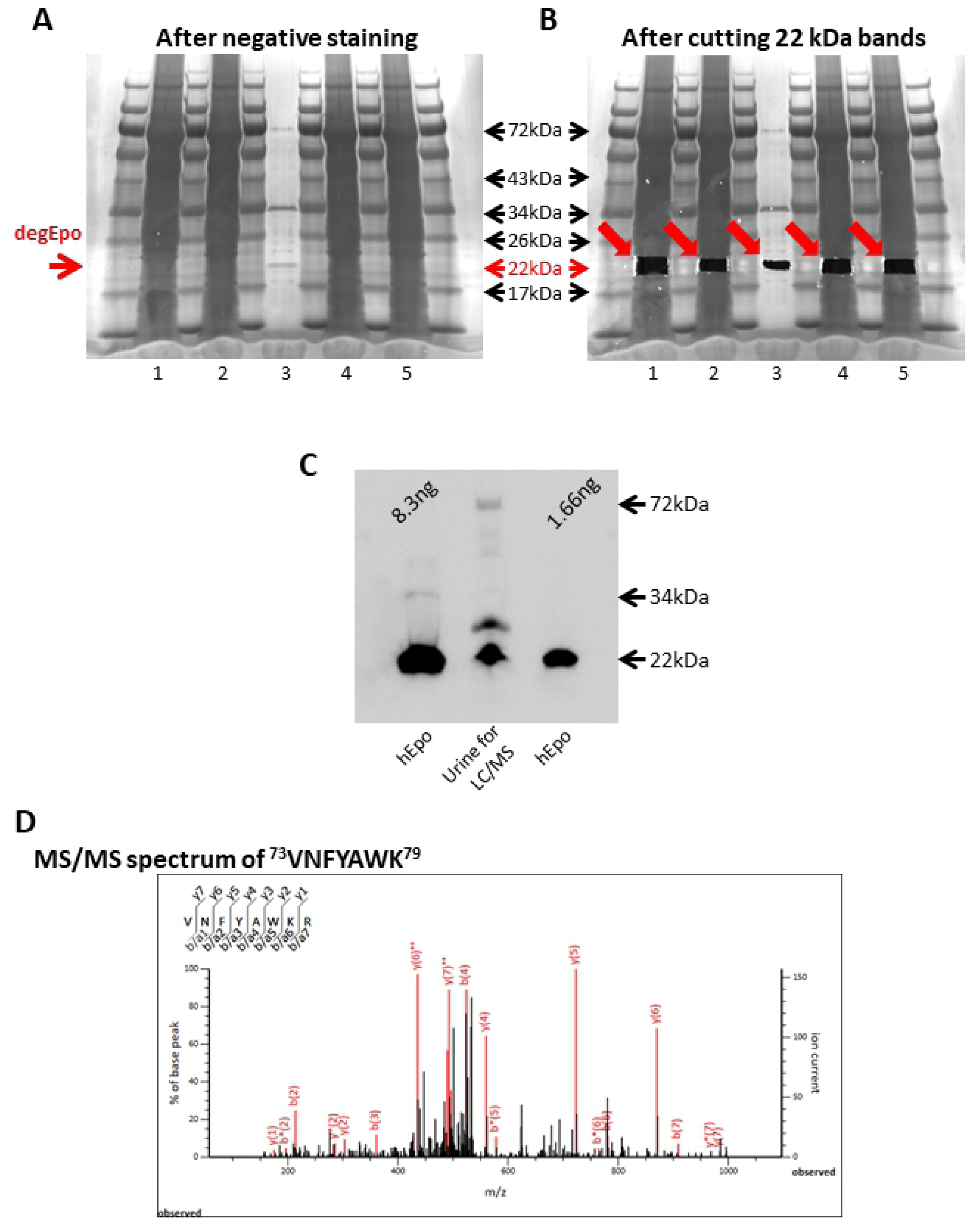
LC/MS detection of recombinant and intrinsic Epo. **A**,**B.** Deglycosylated recombinant human Epo and urine samples of anemic patients were subjected to SDS-PAGE and negative staining. The 22 kDa bands were excised and subjected to LC/MS. Although the recombinant human Epo (8.3 ng) was analysed by negative staining (lane 3), no 22 kDa band was observed in the human urine samples (concentrated from 3.1 ml of urine, lanes 1, 2, 4 and 5). **C.** Western blotting of urine sample used for LC/MS analysis. 12.5 μl of concentrated urine was used in Fig.C and 15μl of concentrated urine was used for Fig. A. **D.** MS/MS spectra of recombinant human Epo peptides: ^73^VNFYAWK^79^. The red line shows the expected peptides, and the black line shows the observed peptides.

### Epo protein expression in hypoxia

Epo mRNA and protein expression in the kidney and liver in hypoxia were examined in rats. HIF1α, HIF2α and Epo mRNA expression in the kidney reached a maximum at 2 hr after hypoxia, and PHD2 mRNA expression in the kidney reached its maximum at 4 hr (Fig. 5A-D). Epo mRNA showed a 200-fold increase in the kidney with no changes in the liver. (Fig. 5A). The plasma Epo concentration showed a 600-fold increase at 4 hr compared with zero time (Fig. 5E). Epo protein expression in the kidney reached its maximum at 4 hr, while the changes in Epo protein expression in the liver were small (Fig. 6A, B). Ususal Western blot showed an approximately 10-fold increase in Epo protein expressions in the kidney, respectively (Fig. 6A, B). Incubation of the kidney samples with deglycosylation buffer without PNGase made the bands clear and the increase of Epo protein expression reached 20-fold increase (Fig. 6C, D). In contrast, deglycosylated Epo protein expression showed an approximately 400-fold increase (Fig. 6C, F), which is very close to the changes in plasma Epo concentration. A very faint band of deglycosylated Epo was observed in the hypoxic liver (Fig. 6E, F).

**Fig. 5.**
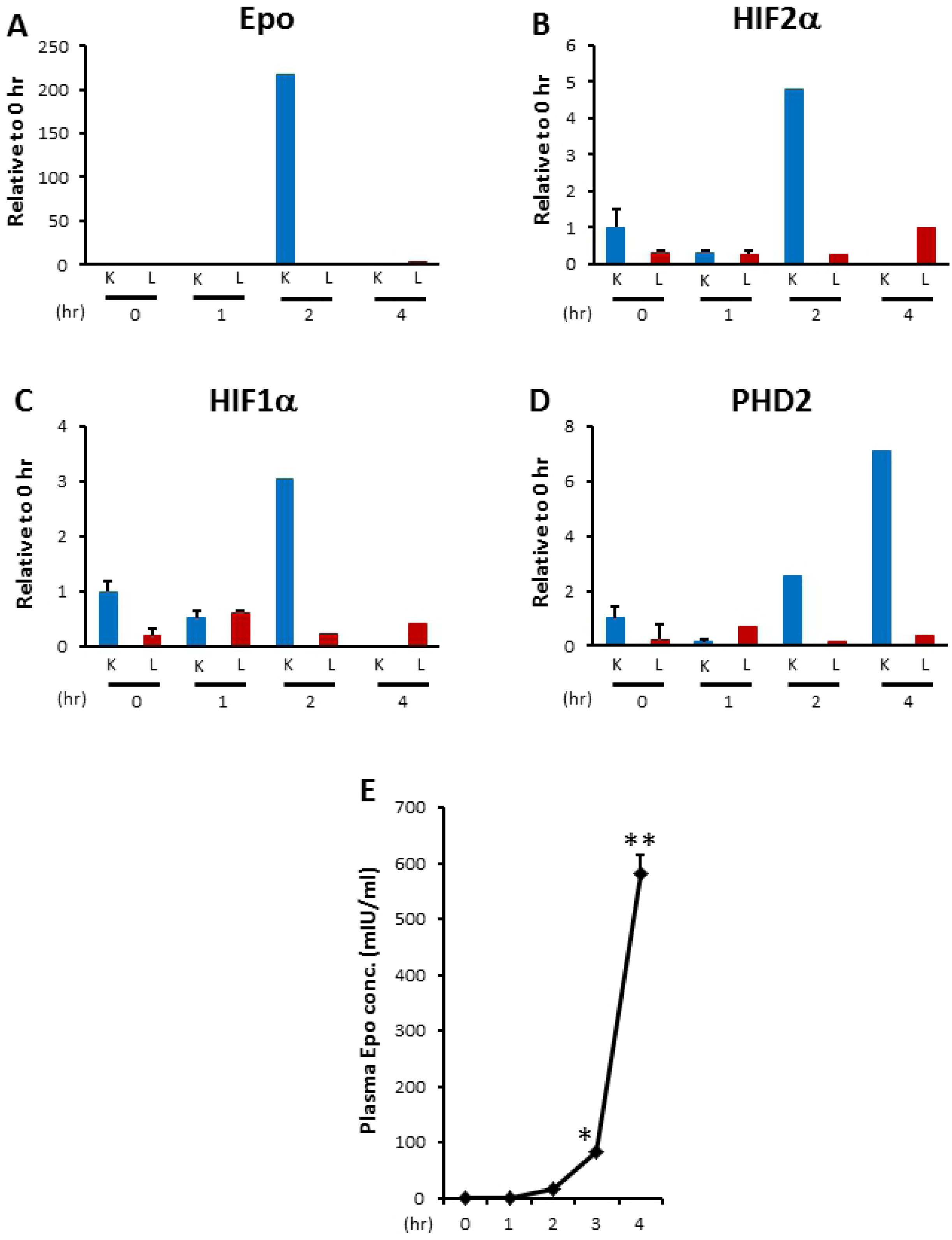
Effects of hypoxia on Epo mRNA expression in the kidney and liver. **A-D.** Effects of severe hypoxia on the mRNA expression of Epo (A), HIF1α (B), HIF2α (C) and PHD2 (D) in the kidney and liver. Severe hypoxia increased HIF1α, HIF2α and PHD2 mRNA expression after 1 hr in the kidney, which was followed by an increase in Epo mRNA expression at 2 hr. All of the above expressions levels decreased thereafter. In contrast, Epo mRNA expression in the liver increased up to 4 hr, which was not related to the changes in HIF1α, HIF2α and PHD2 mRNA expression. n=3-4, K0, K1, K2 and K4; zero time, 1, 2 and 4 hr after the induction of hypoxia in the kidney. L0, L1, L2 and L4: same time course in the liver. **E.** Changes in plasma Epo concentration during severe hypoxia. The plasma Epo concentration significantly increased after 3 hr. n=3-7. * p<0.05, **p<0.001 using ANOVA and multiple comparison with Dunnett’s test.

**Fig. 6.**
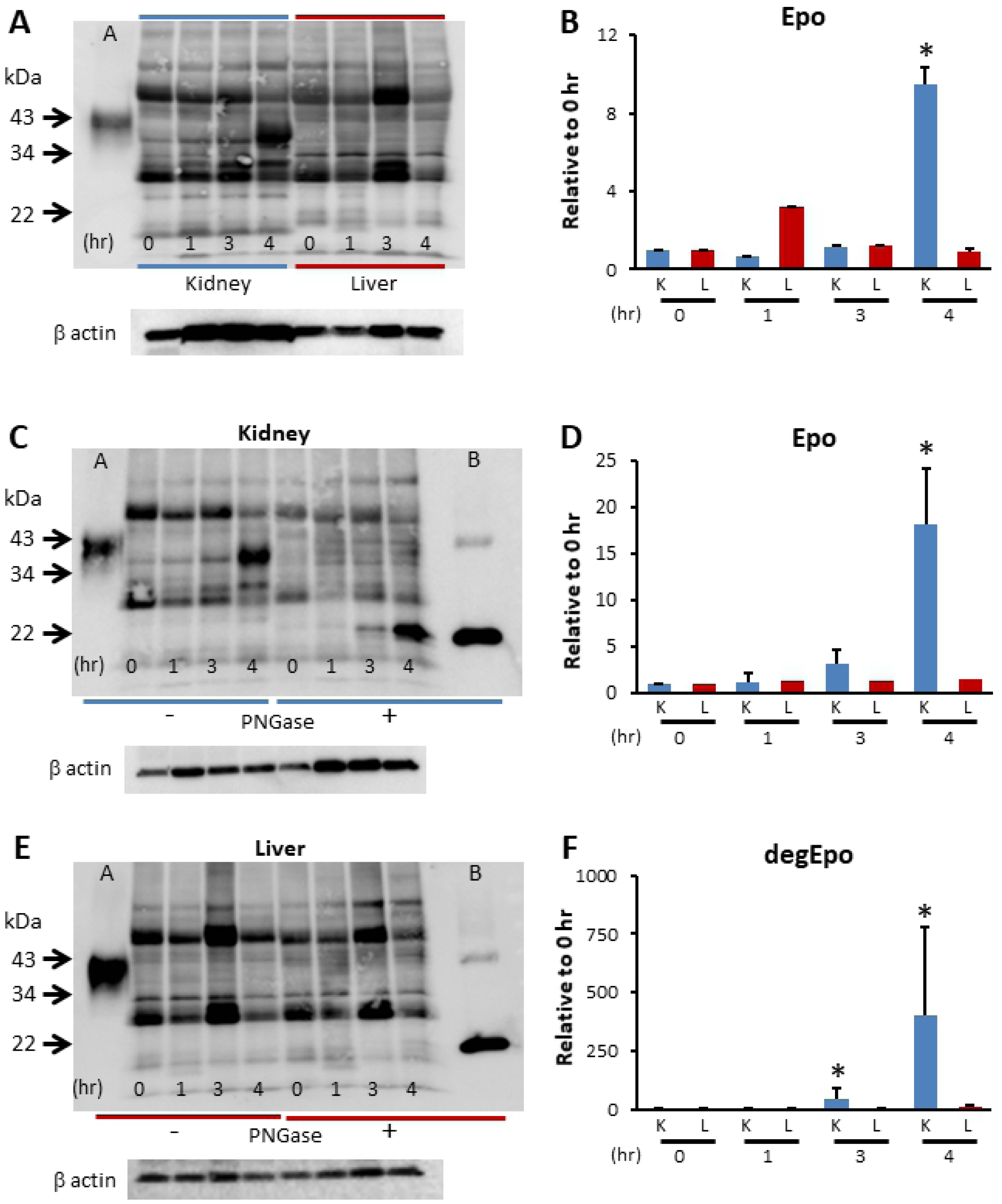
Effects of hypoxia on Epo protein expression in the kidney and liver. **A, C, E.** Western blot analysis of Epo expression in the kidney and liver. A typical gel is shown in Fig. A, C and E, and the analysed data are shown in Fig. B, D and F. Severe hypoxia increased Epo protein expression in the kidney at 4 hr by 10-fold but did not increase Epo protein expression in the liver (A, B). n=4, * p<0.05. Western blot analysis of Epo protein expression after deglycosylation in the kidney (C) and the liver (E). Glycosylated Epo protein expression increased by 20-fold after 4 hr (D). Deglycosylated Epo expression was observed from zero time to 4 hr. The expression increased by 400-fold after 4 hr (F), which was close to the changes in the plasma Epo concentration (Fig. 5E). In contrast, Epo protein expression in liver did not increase under hypoxia (E, F). n=4-6, * p<0.05 using the Kruskal-Wallis test and multiple comparisons by the Shirley-Williams test.

### Immunohistochemical Epo protein expression

Immunohistochemistry showed that renal proximal and distal tubules in the cortex were weakly stained under basal conditions (proximal tubules < thick ascending limbs, distal convoluted tubules) (Fig. 7A, C). Severe hypoxia caused increased Epo staining of the interstitial cells around proximal tubules in the deep cortical area but decreased staining in tubular cells, as in our previous report using in situ hybridization (Fig. 7B, D).

**Fig. 7.**
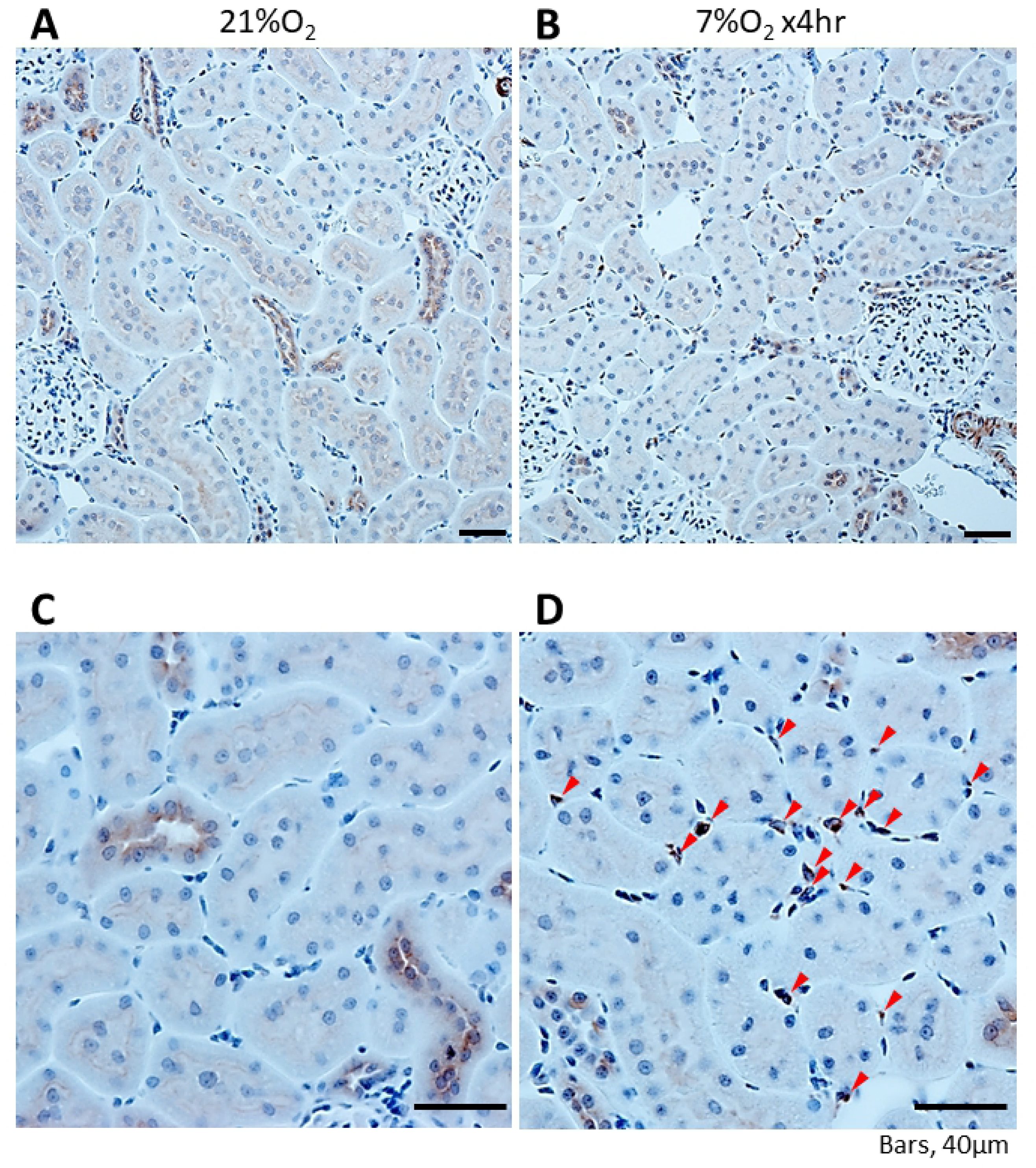
Immunohistochemical analysis of Epo protein expression in the kidney. Epo protein was observed in proximal and distal tubules at 21% O_2_ (A, C). Severe hypoxia (7% O_2_, 4 hr) increased Epo protein expression in the interstitial cells (arrowhead) while slightly decreasing the expression in the tubules (B, D).

## Discussion

We detected Epo protein and ESAs by the combination of usual Western blotting and LS/MS for the first time. Using a new method of Western blotting, we succeeded in the detection of urinary Epo and ESAs. However, intrinsic Epo and ESAs in plasma could not be detected even by our Western blot. The incubation of plasma in deglycosylation buffer resulted in the appearance of bands at 34-43 kDa, and deglycosylation caused a shift of those bands to 22 kDa, except for that of epoetin β pegol (CERA). LC/MS analysis of the 22 kDa band from anemic patient’s urine revealed human Epo. The sensitivity of our Western blotting is higher than that of LC/MS.

One of the findings of our new method is that detection limit of Epo protein is increased by deglycosylation. Detection limit of glycosylated and deglycosylated recombinant human Epo was 370 and 37 pg, respectively (Fig 2B1). The detection limit of deglycosylaed recombinant rat Epo was 3.7 pg (Fig 2B2). Therefore, the deglycosylation increased the detection limit of Eo by 10-100 times. Therefore, accurate quantitative estimates of Epo can be obtained by measuring deglycosylated Epo. Although Epo is detected directly in the urine, the estimation of deglycosylated Epo in the urine would be more accurate.

Our new method will change the tests for Epo doping. Currently, Epo doping is detected by IEF and/or SAR-PAGE or LC/MS after pre-purification of the samples [18, 19]. Our method does not require any pre-purification of the samples. Concentrated urine can be used directly for Western blotting. Blood samples should be deglycosylated to reduce non-specific bands. Intrinsic Epo and ESAs are distinguished simply by band size. To completely confirm the presence of ESAs, cut gels should be checked by LC/MS. More than 1-2 ng of Epo was required to detect Epo by LC/MS, while the detection limit of Epo by our Western blotting is 3.7-37 pg. Since plasma or serum contains a lot of proteins, concentrated plasma becomes very high osmolality and is difficult to use for Western blotting. In contrast, urine has usually no protein except patients with CKD, concentrated urine can be used for Western blotting.

Our new method allowed conclusions regarding unsolved questions about the sites of Epo production in response to severe hypoxia/anemia. Since the increase in Epo production in the kidney was not high enough compared to the changes in plasma Epo concentration and gene expression in the kidney, liver participation has been suggested [2-5, 14]. The difficulty of Epo protein detection by Western blot was the main reason. We showed that deglycylation increased the sensitivity of Epo detection by 10-100 times. Deglycosylated Epo expression showed a 400-fold increase, which is very close to the change of Epo concentration in plasma. Deglycosylated Epo expression in the hypoxic liver was very low. The increases of HIF1α and HIF2α mRNA expression as well as Epo mRNA were observed in the hypoxic kidney but not in the hypoxic liver. The increase of PHD2 mRNA expression and a large decrease of Epo mRNA expression were observed in the kidney 4 hr after hypoxia. HIF2α has a key role for Epo production and PHD2 has a key role for the degradation of Epo [26-28]. These data clearly show that the kidney is the main and sole site of Epo production in response to severe hypoxia. Although plasma Epo is very low in normal rats and humans, control rat kidneys showed deglycosylated Epo production, and immunohistochemistry showed Epo production in the cortical nephrons. Mujais and colleagues reported Epo mRNA expression in renal tubules using microdissected nephron segments in cobalt chloride-injected rats [29]. We have previously shown that fludrocortisone stimulated Epo production by the intercalated cells of the collecting ducts [20]. Our immunohistochemistry also showed that kidney interstitial cells respond to severe hypoxia by producing Epo. Yamamoto and colleagues showed that the site of Epo production by severe anemia is the interstitial cells using EPO promoter-driven GFP expression [8, 13]. Since 27 kDa GFP goes into nucleus, they may have overestimated the role of Epo production by interstitial cells in severe anemia. Since the cytoplasm of interstitial cells is very pale, Epo production by interstitial cells under hypoxia may not be as strong as expected. These data show that kidney nephrons produce Epo under control conditions and that kidney interstitial cells produce Epo in response to severe hypoxia or anaemia.

In conclusion, our data showed that Epo protein can be detected in urine and tissue samples by direct Western blot analysis and in blood after deglycosylation. Our data also showed that the kidneys have dual Epo production systems, low production by the nephron under normal conditions and hypoxia or anemia-induced high production by the interstitial fibroblast-like cells, and that the kidney is the main and sole site of Epo production in response to hypoxia or anaemia. Our method will fundamentally change Epo doping and detection.

## Acknowledgements

Our manuscript was edited for proper English language by NPG Language Editing Service (4221-D9DA-8D07-E1B1-3D9P).

## Author contributions

YY, YI, KK and HN designed the research; YY, YN, HI, YoS, YN, and HN performed the animal research; YI, TF, KY, TU, and HN performed western blot analysis; YY, TO, YuS, and KK performed IHC, TF, TaY, NK and HN performed RNA extraction and PCR; YI and HN performed the statistical analyses; and TeY performed LC/MS. MM and YuS advised on the experimental design and data interpretation.

## Additional information

Correspondence and requests for materials should be addressed to.

